# The causal role of the inferior temporal cortex in visual perception

**DOI:** 10.1101/2022.10.24.513337

**Authors:** Elia Shahbazi, Timothy Ma, Martin Pernuš, Walter Scheirer, Arash Afraz

## Abstract

Neurons in the inferotemporal (IT) cortex respond selectively to complex visual features, implying their role in object perception. However, perception is subjective and cannot be read out from neural responses; thus, bridging the causal gap between neural activity and perception demands independent characterization of perception. Historically though, the complexity of the perceptual alterations induced by artificial stimulation of IT cortex has rendered them impossible to quantify. Here we addressed this old problem by combining machine learning with high-throughput behavioral optogenetics in macaque monkeys. In closed-loop experiments, we generated complex and highly specific images that the animal could not discriminate from the state of being cortically stimulated. These images, named “perceptograms” for the first time, reveal and depict the contents of the complex hallucinatory percepts induced by local neural perturbation in IT cortex. Furthermore, we found that the nature and magnitude of these hallucinations highly depend on concurrent visual input, stimulation location, and intensity. Objective characterization of stimulation-induced perceptual events opens the door to developing a mechanistic theory of visual perception. Further, it enables us to make better visual prosthetic devices and gain a greater understanding of visual hallucinations in mental disorders.

**One-Sentence Summary:** Combining state-of-the-art AI with high-throughput closed-loop brain stimulation experiments, for the first time, we took “pictures” of the complex and subjective visual hallucinations induced by local stimulation in the inferior temporal cortex, a cortical area associated with object recognition.

---

Artificial stimulation of neurons in high-level visual cortical areas induces complex hallucinatory visual percepts (*3–6*). Scientific characterization of these visual percepts poses a serious methodological challenge due to their complex and subjective nature, yet it has inspired a multigenerational effort in systems neuroscience as it bridges the causal gap between patterns of neuronal activity in the brain and elements of visual perception (*1, 7, 8*). From a translational point of view, understanding the causal underpinnings of visual hallucinations induced by local brain stimulation is necessary to develop prosthetic devices that restore vision by direct brain stimulation (*9, 10*). This knowledge also shapes the building blocks for understanding visual hallucinations in mental disorders and altered states of consciousness (*11–13*).

## Perceptography: concept and methodology

In this study, we created a machine learning structure and used it in combination with high-throughput behavioral optogenetics in macaque monkeys in order to, for the first time, produce pictorial descriptions of the perceptual events induced by brain stimulation in the high-level visual cortex. These pictorial descriptions, called *perceptograms*, provide unbiased and parametric yet rich accounts of the visual perceptual events following optogenetic activation of ∼1mm3 neural subpopulations in the inferior temporal (IT) cortex. The basic idea behind our quest was simple: guided by the animals’ behavior, is it possible to evolve specific image perturbations that resemble the sense of being stimulated in a given cortical locus in the absence of physical stimulation?

We performed viral injections in the central IT cortex of two macaque monkeys (*Macaca mulatta*) in order to express the excitatory opsin C1V1 under the CaMKIIa promoter in a ∼5×5mm area of the cortex. We then implanted arrays of LEDs (Opto-Array) on the virally transduced cortical area as well as the corresponding position in the opposite hemisphere where no viral injection was performed. The Opto-Array allows safe, rapidly reversible, and high-throughput optical stimulation of ∼1mm3 subregions of the targeted cortex, although it doesn’t allow neural recordings. Technical details about the Opto-Array and relevant surgical protocols can be found in our earlier reports (*2, 6*).

The two monkeys were trained to detect and report a brief optogenetic stimulation impulse delivered to their IT cortex while fixating on a short video (1 second) created by a generative adversarial neural network (GAN) (Fig 1.a). It has been previously shown that monkeys can easily learn this simple task (*14, 15*), which remained the sole task expected from the animals throughout the study. Our earlier results suggest that the animals perform this task (in the IT cortex) using the visual events induced by cortical stimulation (*6*). The animals initiated each trial by holding fixation on a central target for 500ms. Then, a natural-looking GAN-generated image was shown for 400ms (seed image) on a gray background. The image subtended 8ox8o of visual angle, and the animals were required to hold fixation at its center throughout the trial. Next and in all trials, the seed image would turn into a randomly perturbed version of itself for 150ms, then turn back into the original image and stay changeless for 450ms. In half of the trials, randomly selected, at the time of image perturbation, an LED was activated on the animals’ IT cortex for 150ms, typically at 3mW photometric power. After the video clip (seed-perturbed-seed), the screen was cleared, and two response targets appeared on the vertical midline (white, 0.4° diameter, 5° above and below the center). The animals then made a saccade to one of the two targets in order to indicate if the trial included a brain stimulation impulse (chance level 50%). The response targets then disappeared, and the animals received a liquid reward for correct responses and a 3.5s timeout for incorrect responses. Trials with broken fixation or latency greater than 3s for making a response were aborted and discarded. These trials were injected into the future stream of trials in a pseudorandom order.

**Fig 1.**
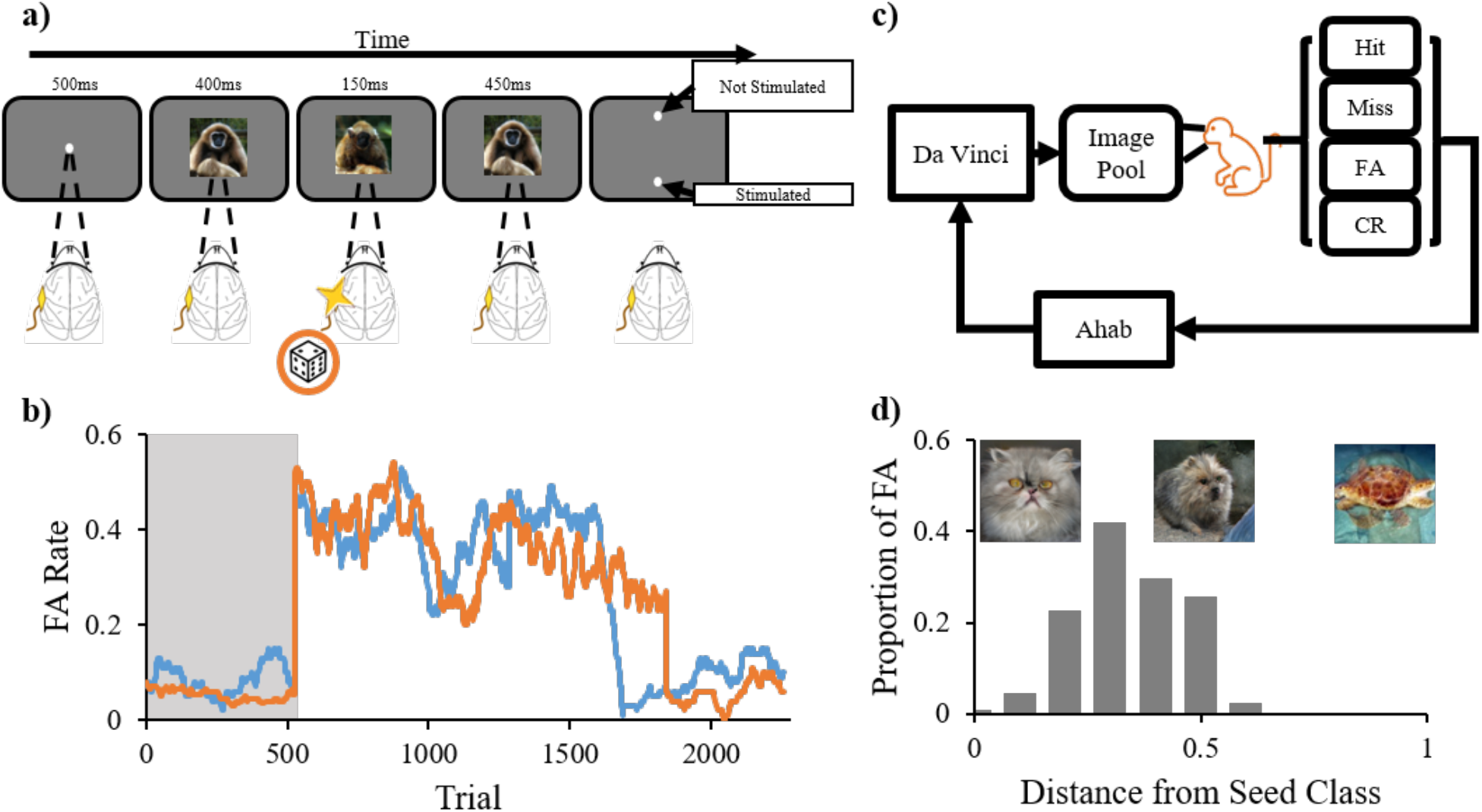
Perceptography paradigm and pipeline. **a)** Cortical perturbation detection (CPD) task. After fixation, a short movie consisting of a 400ms presentation of a seed image followed by 150ms of a perturbed image, and then 450ms of the original seed image was played. In 50% of trials at random, an ∼1mm^3^ locus in the IT cortex was optogenetically stimulated for 150ms at the same time as the perturbed image presentation. The animals were rewarded for correctly identifying if trials contained brain stimulation or not. **b)** The first training day with dynamic stimuli. The abscissa shows trials, and the ordinate represents the false alarm rate. The shaded region indicates trials performed while fixating on a solid static image (initial training for the task). The rest of the plot shows the FA rate after the 150ms image perturbation was first introduced to the training regime. Both animals first took the image alterations as “stimulated trials” at very high rates but learned within a few hundred trials to ignore most of the image perturbations and veridically detect the cortical stimulation. Blue: Monkey Sp, Orange: Monkey Ph. **c)** Perceptography pipeline. The illustrator engine, DaVinci, generated a pool of randomly perturbed images. The optimizer engine, Ahab, analyzed the monkeys’ performance in the CPD task to extract the image features that increased the likelihood of behavioral false alarms. Ahab sent the optimized parameters to DaVinci to generate new pseudo-random image perturbations. These Ahab-optimized images were heavily diluted with random DaVinci images and injected back into the image pool for the next cycle of perceptography. **d)** Proportion of behavioral false alarms as a function of the magnitude of image perturbation. The abscissa shows the normalized feature distance of each randomly perturbed image from its seed image. The ordinate represents the behavioral FA rate for the first pool of DaVinci images. Images above the chart are examples of the visual change corresponding to the distances of 0, 0.5, and 1 from the seed class, respectively, from left to right.

As reported in our earlier work, the animals learned to perform the cortical perturbation detection (CPD) task quickly while fixating on static images (without any image perturbation), and they were not able to detect cortical illumination over the intact cortical area where no viral injection was performed (*6*). On the first day, when the dynamic image perturbation in the middle of the trial was introduced, both animals mistakenly mixed the image perturbations with cortical stimulation, and as a result, their false alarm (FA) rate dramatically increased from 8 and 5.2 percent to 39.2 and 37.6 percent, respectively for monkeys Sp and Ph. This strongly suggests that optogenetic stimulation of the IT cortex induces a “visual” perturbation that can be mixed up with an image perturbation on the screen. Note that the FAs are the trials in which no cortical stimulation was delivered, yet the animal reported the trial as “stimulated.” Also, note that a FA is considered a behavioral mistake and is never rewarded together with the Miss trials when a stimulated trial is reported as non-stimulated. Within a single day, both animals learned to discriminate IT stimulation from the image perturbations on the screen and performed the task with high performance at 90.2 and 89 percent correct and only 8.3 and 6.2 FA rates (respectively for Sp and Ph). This remarkable observation is documented in Fig 1.b.

## Results

### The state of brain stimulation can be mimicked by images

After the training phase and once stable high-performance levels (above 80% for both animals) were achieved, the animals entered the first phase of behavioral data collection. While the monkeys performed the simple CPD task for tens of thousands of trials, under the hood, two learning systems controlled the experiment with the goal of evolving specific image perturbations that increase the chance of behavioral false alarms. We refer to these two systems as DaVinci and Ahab (see Fig 1.c). DaVinci is our image illustrator engine, a structure powered by BigGAN trained on the ImageNet dataset (*6, 16*). DaVinci was tasked with creating multiple random image mutations for each seed image (see Methods). Ahab is our feature optimizer (see Methods) tasked with tracking the animals’ behavioral responses to DaVinci’s random image perturbations. Ahab learned from the animals’ behavioral mistakes and gave DaVinci feedback to produce image perturbations that would increase the FA rate. An increase in the FA rate (trials without stimulation reported as stimulated) could result from a general increase in task difficulty, which would also increase the Miss rate (trials with stimulation reported as non-stimulated). To avoid this, Ahab was set to aim at specifically increasing the FA rate without changing the Miss rate (see Methods).

The image evolution process started with 5-6 image seeds, for each of which DaVinci created 400-1000 randomly perturbed images. Each of these image perturbations was presented to the animal at least five times in the course of multiple days (a total of 10K-30K behavioral trials). While image perturbations are done randomly over a nearly infinite feature space (see Methods), the amplitude of these perturbations can vary: small perturbations randomly but subtly change the image, while large perturbations induce random yet massive pictorial alterations. Fig 1.d. plots the behavioral FA rate as a function of image perturbation magnitude. We found this result encouraging as it shows that the behavioral false alarm rate in the CPD task can be systematically manipulated by altering the image. It also shows that increasing the magnitude of image perturbation does not monotonously increase the chances of a non-stimulated trial being taken as stimulated (FA). The distribution of behavioral false alarms over image alterations of various sizes reflects the magnitude of the perceptual perturbations induced by cortical illumination for the stimulation intensity used in this experiment (3mW).

### Artificial intelligence learns from the brain how to trick it

Next, Ahab scored each image perturbation and selected the ones that induced a higher FA rate without increasing the Miss rate (see Methods). Ahab guided DaVinci to create an image family for each surviving image, including the original image and 2-6 mutated children. These image families were then presented to the animals in the context of the next round of behavioral testing, and the images that were scored high by Ahab received the chance to mutate again and make their own children. This process was repeated until at least one of the image families passed the threshold of 60% FA over at least 12 presentations. This typically took five iterations of the entire process, involving 1-5K Ahab-optimized image presentations. The image that scored highest within a winning family was named a *perceptogram*, as viewing it was hard for the animal to distinguish from the perceptual state induced by brain stimulation. The entire process was accordingly coined: *Perceptography*. Throughout the course of each round of perceptography, a single LED of the Opto-Array was selected and used. The intensity of the LED was adjusted at each new cortical position in order to keep the behavioral output under ceiling performance.

The process of Perceptography, if successful, would increase the FA rate across generations of images. This could untrain the animals over the course of time because we only reward objectively correct choices. To avoid untraining the animals by this procedure, we heavily diluted Ahab-optimized image families with non-optimized DaVinci images as the evolution progressed (50 to 80 percent non-optimized). While the optimized images were heavily diluted by DaVinci images, the animal’s FA rate kept increasing specifically for those images as the evolution progressed. Fig 2.a shows the monkeys’ FA rate as a function of session number for DaVinci and Ahab optimized image families. As shown in the figure, the FA rate remained at a constant level of 2.8-4.1% and 4.1-6.1% (respectively for Sp and Ph) for DaVinci images, but Ahab optimized image families induced more FAs increasingly as the process unfolded. Fig 2.b and 2.c show the evolution process for a typical perceptogram starting from a large variety of image perturbations and converging to a specific one.

**Fig 2.**
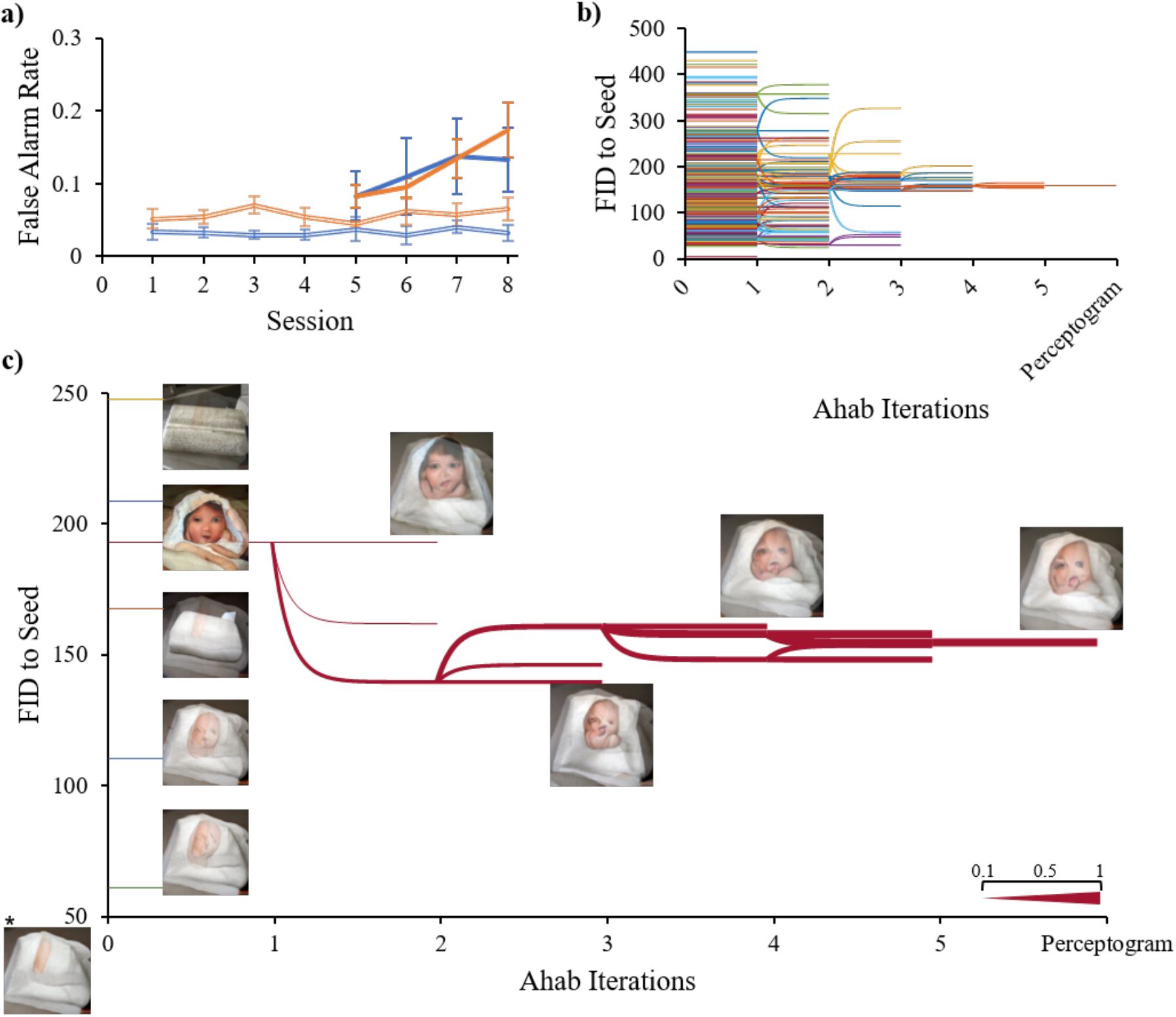
Evolution of perceptograms. **a)** The false alarm rate for non-optimized (Davinci, double lines) and optimized (Ahab, solid lines) perturbed images. The abscissa indicates the progress of perceptography across sessions. The ordinate shows the FA rate. Blue: Sp, Orange: Ph. Ahab optimized images induced a significantly higher false alarm rate (df = 74 and 76, p = 0.025 and 0.002 for Sp and Ph, respectively, Welch’s t-test). Error bars indicate ±1 standard error of mean **b)** Evolution dendrogram. Each colored line represents a single image family. To survive the iterations of Ahab optimization, image families had to maintain a cumulative false alarm rate of over 50%. The ordinate shows the Fréchet Inception Distance (FID) between each perturbed image and its corresponding seed image. The abscissa shows iterations of the perceptography procedure. **c)** Example of a perceptogram image family tree. The abscissa and ordinate are the same as in subplot b. The legend on the bottom right shows how the thickness of each branch corresponds to its false alarm rate. 5 examples of image mutations from the initial DaVinci pool are shown together with the winning image family tree. The asterisk indicates the seed image.

### Robustness of the results

Is it possible that some image perturbations survive through the pipeline by chance without being meaningful to the animals? We bootstrapped the data, but instead of letting the animals determine the distribution of FAs for each trial, we distributed them randomly. Fig 3.a shows the results. If the false alarms were randomly distributed across image presentations, the best image family would have a cumulative false alarm rate significantly lower than the image families selected by the perceptography process. More interestingly, data shows that the contents of these behaviorally selected images are related. In fact, as Fig 3.b shows, the images selected independently by the animals’ behavior across families share increasingly more features as the process continues. These analyses show that using the image evolution process presented here, it is extremely unlikely to get an image tagged as a perceptogram just by chance. They also show that the image evolution process is not a random stray trajectory; instead, it is systematically guided by the animals’ choices and converges on specific answers. Despite these statistical encouragements and in order to fully cross-validate these findings with a fresh set of data, we performed the entire perceptography procedure on the same image seed, cortical position, and stimulation intensity once again for each animal. Fig 3.c shows how two independent perceptography procedures converged on similar answers. These procedures, each lasting ∼17 work days, were performed 24 and 10 days apart from each other in monkeys Sp and Ph, suggesting that the perceptual effects of repeated optogenetic stimulation in a given cortical position remain stable at least over the course of ∼ one month.

**Fig 3.**
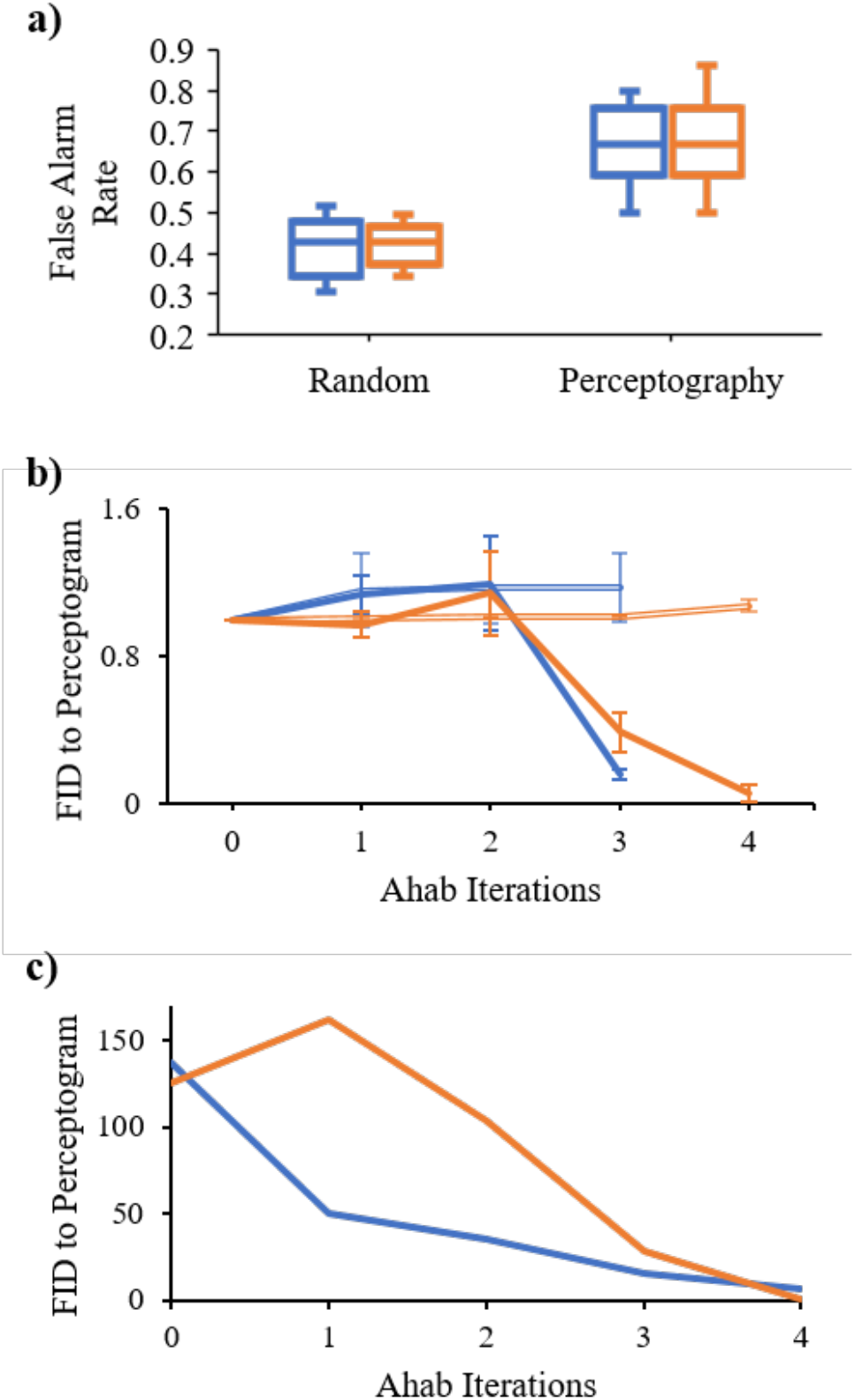
Evolution trajectory of perceptograms; random or guided? **a)** High FA rates cannot be achieved by random selection of image families across iterations. Left: the distribution of maximum false alarm rates achievable in bootstrapped data where FA scores are randomly assigned to images at each iteration of perceptography. Perceptograms, images that evolved guided by the animals’ behavior, had significantly higher FA rates compared to the best images produced by the bootstrapping procedure (df = 13 and 17 for Sp and Ph, respectively, p<0.0001 for both, Welch’s t-test). Right: the distribution of perceptogram false alarm rates. Blue: Sp, Orange: Ph. Error bars indicate the minimum and maximum rates **b)** Convergence to similar images across Ahab iterations. The abscissa represents Ahab’s iterations of optimization. The ordinate shows the FID feature distance. The solid lines represent the FID distance of the final perceptogram from images in each optimization iteration, excluding the perceptogram family. Independently optimized images get more similar to each other, and the final perceptogram as the process unfolds. Double lines represent the same for the bootstrapped data where family survival is randomly chosen. As the image pool was optimized by Ahab, the distance (FID) between the optimized pool and the to-be-discovered perceptogram decreased. Note that the images were not selected for similarity but based on the behavioral FA rate they evoke (Blue: Sp, Orange: Ph). Error bars indicate ±1 standard error of mean **c)** Independent evolution of similar perceptograms. Two independent rounds of perceptography were performed for each monkey (Blue: Sp, Orange: Ph). The axes are similar to the subplot b. The line plot shows the FID distance of the optimized images of the second round of perceptography with the perceptogram obtained from the first round.

A design feature of the experiments reported here is that we use the same image perturbations in both stimulated and non-stimulated trials. This balancing feature is crucial in order to take away all potential image cues and forces the animals to read out only the cortical stimulation for performing the task. This feature, however, introduces a measurement uncertainty to the process. As a result of stimulus balancing, all stimulated trials include two perturbation components, one is on the screen, and the other comes from the brain stimulation. The screen component is not informative and varies at every trial, and the monkey is incentivized to ignore it and detect the cortical component. Now, when the presentation of a perceptogram in a non-stimulated trial elicits a behavioral FA, the animal matches the perceptogram to the net perceptual effect of cortical stimulation (constant across stimulated trials) plus a baseline random non-informative component (variable across trials). While this introduces an inherent uncertainty in the procedure, in that the measurement process affects the measure of interest, since the image perturbations are mostly small and random, their net effect is not expected to drift far from the original seed image. From the point of view of an incentivized observer, most image perturbations are expected to be perceived as irrelevant, except the ones that warp the seed image in the same direction as induced by the brain stimulation. If true, this would increase the chance of reporting the trial as stimulated in both stimulated and non-stimulated conditions. Fig 4.a shows that the hit rate is higher than baseline when the cortex is stimulated while looking at perceptograms. Although, given the high baseline hit rate, the reward that the monkey gains at stimulated trials (grand average 7.9% and 6.5% for Sp and Ph) are far less than the reward loss at non-stimulated trials when perceptograms are presented (grand average 60.0% and 64.3% for Sp and Ph). Moreover, it seems that the animals psychophysically rely on contrasting stimulation with the solid seed images presented before and after the stimulation more than the perturbed image. In an experiment, we showed image perturbations of one seed image (150ms) temporally sandwiched between images of another seed. This was done for two seed images in each monkey. The FA rate dramatically decreased in all cases, indicating that the perceptual effect of stimulation is perceived and matched by the animals mainly in temporal contrast to the seed image. Specifically, the false alarm rate dropped to 0% and 2% (out of 50 presentations) in Sp and Ph, respectively.

**Fig 4.**
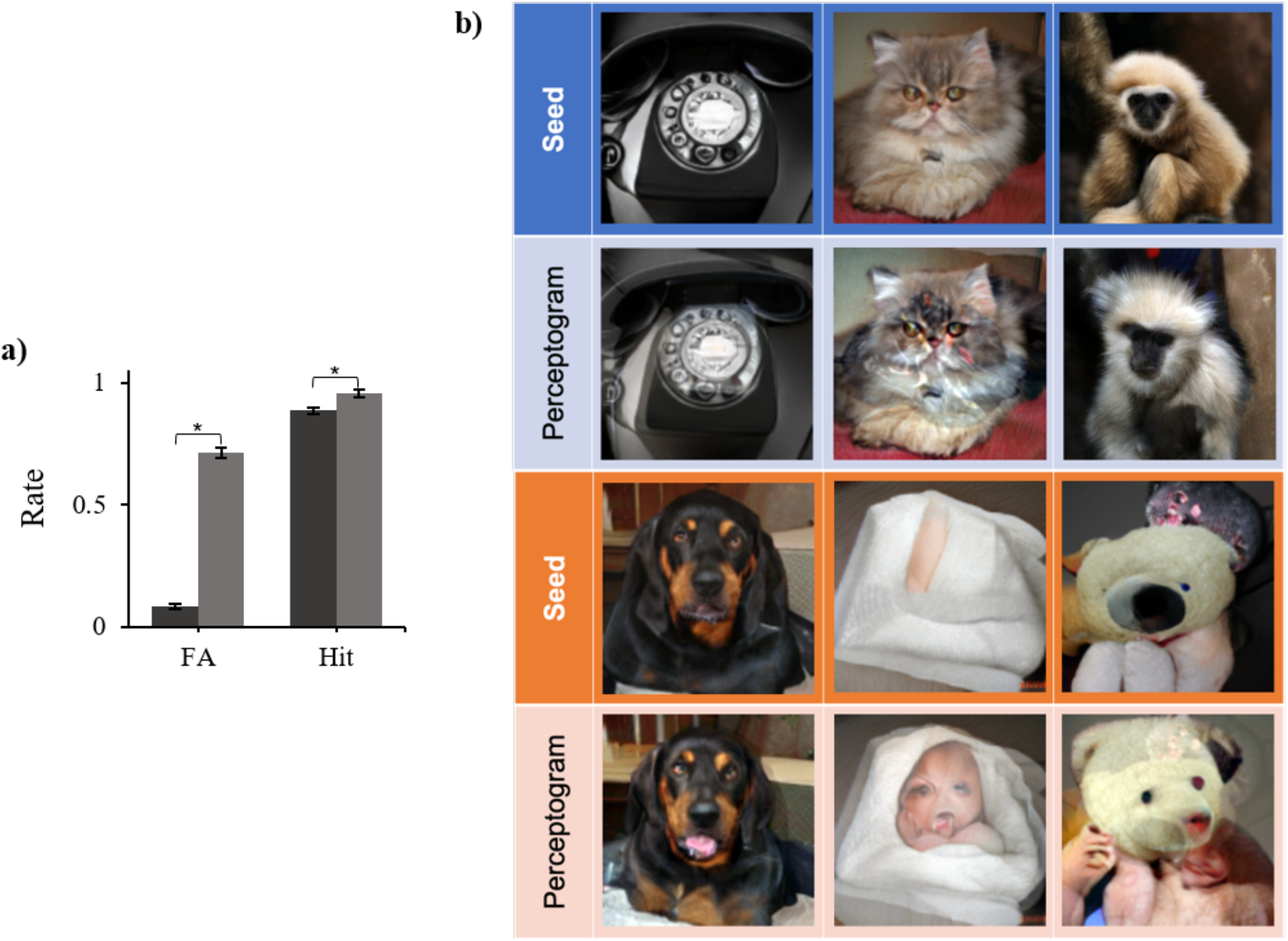
The effect on hit rate and some examples of perceptograms. **a)** Perceptograms increase the hits as well as FAs. The false alarm rate evoked by the perceptograms (light gray) is significantly higher than that of the non-optimized DaVinci image pool (dark gray) (df = 30, p <0.001). The hit rate is also significantly higher in perceptograms even though the effect is smaller due to a ceiling effect (df = 30, p < 0.001). Error bars indicate ±1 standard error of mean **b)** Examples. Three examples are shown from each monkey (Blue shades: Sp, Orange shades: Ph); in each block, the top row indicates the seed images, and the bottom row shows their corresponding perceptograms.

### Effects of stimulation intensity

Fig 4.b shows examples of perceptograms obtained from the two animals. As an independent sanity check, we hypothesized that if a perceptogram truly reflects the perceptual changes induced by cortical stimulation, the magnitude of image perturbation in the winning perceptograms should increase if we increase the cortical stimulation intensity. To test this, we performed independent perceptography procedures on similar cortical positions, each at two different cortical illumination powers. Fig 5.a and 5.b demonstrate that the amount of feature warping in the winning perceptograms was remarkably higher when higher cortical illumination was applied. Examples of perceptograms at each level of cortical illumination are shown in Fig 5.c. Consistently, the baseline miss rate of both animals was slightly but significantly lower in the high illumination condition, as shown in Fig 5.d.

**Fig 5.**
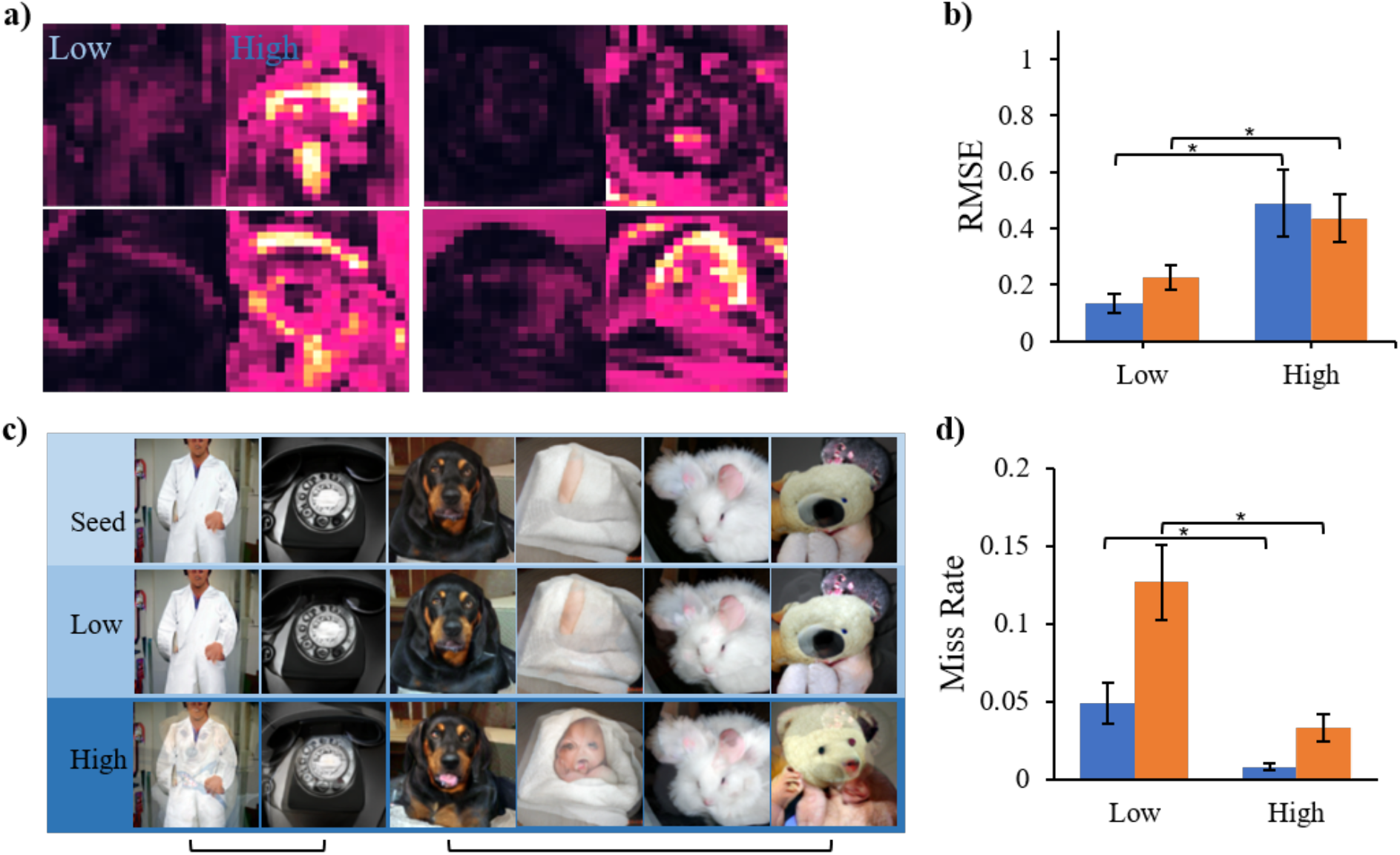
Effects of stimulation intensity. **a)** Examples of heat maps depicting the image changes between the perceptograms and their corresponding seeds. In each block of heatmaps (left Sp, right Ph), the left column includes the perceptograms obtained from low illumination perceptography. In contrast, the right column depicts the high illumination perceptograms in the same LED channel and monkey. **b)** More intense cortical illumination warps the resulting perceptograms further away from their seed images. In both animals, the perceptograms obtained with higher intensity of stimulation had significantly higher distances from their image seeds compared to the perceptograms resulting from low-intensity cortical illumination (Root Mean Squared Error (RMSE): df = 8 and 13, p = 0.025 and 0.026 for Sp and Ph respectively, Welch’s t-test). Blue: Sp, Orange: Ph. Error bars indicate ±1 standard error of mean **c)** Examples of perceptograms obtained with low and high stimulation intensities. Top: seed images. Middle: perceptograms obtained with low cortical illumination. Bottom: perceptograms obtained with higher illumination power. The brackets under the subplot indicate the perceptograms obtained from the same channel **d)** The effect on the behavioral miss rate. Increasing the illumination intensity of the LEDs significantly decreased the behavioral miss rate in both monkeys (df = 13 and 8, p = 0.026 and 0.023 for Sp and Ph, respectively, Welch’s t-test). Blue: Sp, Orange: Ph. Error bars indicate ±1 standard error of mean.

### Perceptograms and natural image manifolds

Overall, the development of each perceptogram cost ∼30-50K behavioral trials collected in the course of 14-20 work days. We performed a total of 32 complete rounds of perceptography over seven cortical locations (3 and 4 for monkeys Sp and Ph), 15 seed images, and six stimulation intensity levels. These results provide pictorial evidence of the visual perceptual hallucinations induced by stimulation of the high-level visual cortex. Examples of a few perceptograms are shown in Fig 4.b. These results show that it is possible to behaviorally exchange the state of local brain stimulation in IT cortex with the state of viewing an image. The similarity of the two states is close enough to make the animals choose to tag non-stimulated perceptogram trials as stimulated, even at the cost of losing reward. While an “ideal perceptogram” is expected to induce a 100% FA rate, the ones found in this study (mean FA rate 70.2%, Median=71%, StD=12) are surprisingly close, given the very low baseline FA rates. The residual from 100% can be due to the imperfection of our image generation engine and/or potential effects of stimulation that are impossible to mimic on a 2D screen (e.g., 3D hallucinations, nonvisual feelings, etc.). Such effects, even if existing, must be very subtle in amplitude because the animals are incentivized to use any clue to receive a reward.

A feature that is expected to emerge from the examination of perceptograms is a common visual element in perceptograms obtained from the same channel. IT cortex is known for its strong object selectivity at the single cell (*17, 18*) as well as ∼1mm3 tissue scale (*19–21*). While the current OptoArray technology doesn’t allow neural recording, rendering us blind with respect to the object selectivity profile of the stimulated neurons, it is reasonable to assume heterogeneity of selectivity at the spatial scale perturbed by a single LED(*6, 19*) in that the perturbed neural population conserves visual selectivity for “a” part of the shape space. Assuming this, one might expect that stimulation of a given site in IT cortex induces the perception of the preferred features of the targeted neurons independent of what is on the screen. The results, though, show a completely different picture. Fig 5.c depicts examples of perceptograms obtained from one cortical position in each of the two monkeys along with their corresponding seed images (more examples are provided in the supplementary materials, Fig S1). The first observation that is apparent in these perceptograms is that their structure strongly depends on the seed image. Perceptograms that come from stimulation of a single point in the IT cortex are typically very different from each other, lacking at least an obvious explicit common visual element. This is consistent with the recent findings about the vast activity landscape of IT neurons(*22*) and the idea that the activity of a neural unit is interpreted by the rest of the brain only in the context of the state of other similar neural units (*8*). These findings strongly encourage recording the neural activity together with perceptography, a point that is further dissected in the *conclusion*.

What do perceptograms reveal about the perceptual effects of brain stimulation in area IT? Inspecting the set of perceptograms produced in our experiments (Fig S1), one first notices that most of the perceptograms show image changes that are off the manifold of natural objects. However, a few seem suspiciously natural; for example, a dog seed image (Fig 4.b bottom block) has turned into exactly the same dog sticking out its tongue, or a monkey (Fig 4.b top block) has turned into a very similar monkey with long light-colored hair and the head turned a few degrees. Consistent with this observation, a scoring algorithm based on Yolov3, a real-time object detection system(*23*) scored 15% (3 out of 20) of the perceptograms as “natural images” (defined as less than 10% change in the main label confidence compared to the seed without introducing any new label with confidence more than 20%). This shows that perturbing the neural activity in ∼1mm3 of the IT cortex pushes the neural state off its natural manifold on most occasions; however, in some cases, the pattern of activity induced by the external stimulus is so that the same neural perturbation creates a naturally meaningful change. Determining when a perturbation lands on the natural manifold of neural activity is a critical step for breaking the code that maps the neuronal activity to perception(*8*).

### Effect of cortical position

Another point that pops out while comparing the perceptograms coming from different LED channels is that anterior channels seem to have induced more holistic changes in the image. While perceptograms express significant pixel deviations from their seed images all along the posterior-anterior axis, the quality of these changes varies systematically. Inspecting the examples presented in Fig 6.a, one would notice that stimulation in the posterior channels of the array distorts the perceived image by adding irrelevant visual features to the contents of perception. However, the anterior channels induce perceptual changes that are identity-preserving and stay on the manifold of natural images. These subjective evaluations can be tested by state-of-the-art object classification tools. Analysis of images shows that perceptograms of the anterior channels of the array tend to retain general features of the seed image as shown by the high confidence in image classification and a low FID distance to the seed (measuring Frechet Inception Distance score, calculates feature vector distance between generated and real images, or any two sets of generated images, Fig 6.b, 6.c. and 6.d.). In contrast, perceptograms of the posterior LEDs express the opposite effect, where additional features are introduced, thus lowering the confidence in image classification and increasing the FID to the seed. Consistent with numerous studies of IT cortex that show a tendency for neural responses to more holistic features along the posterior-anterior axis of the cortex (*8, 19, 24–27*) this finding supports the causality of the relationship.

**Fig 6.**
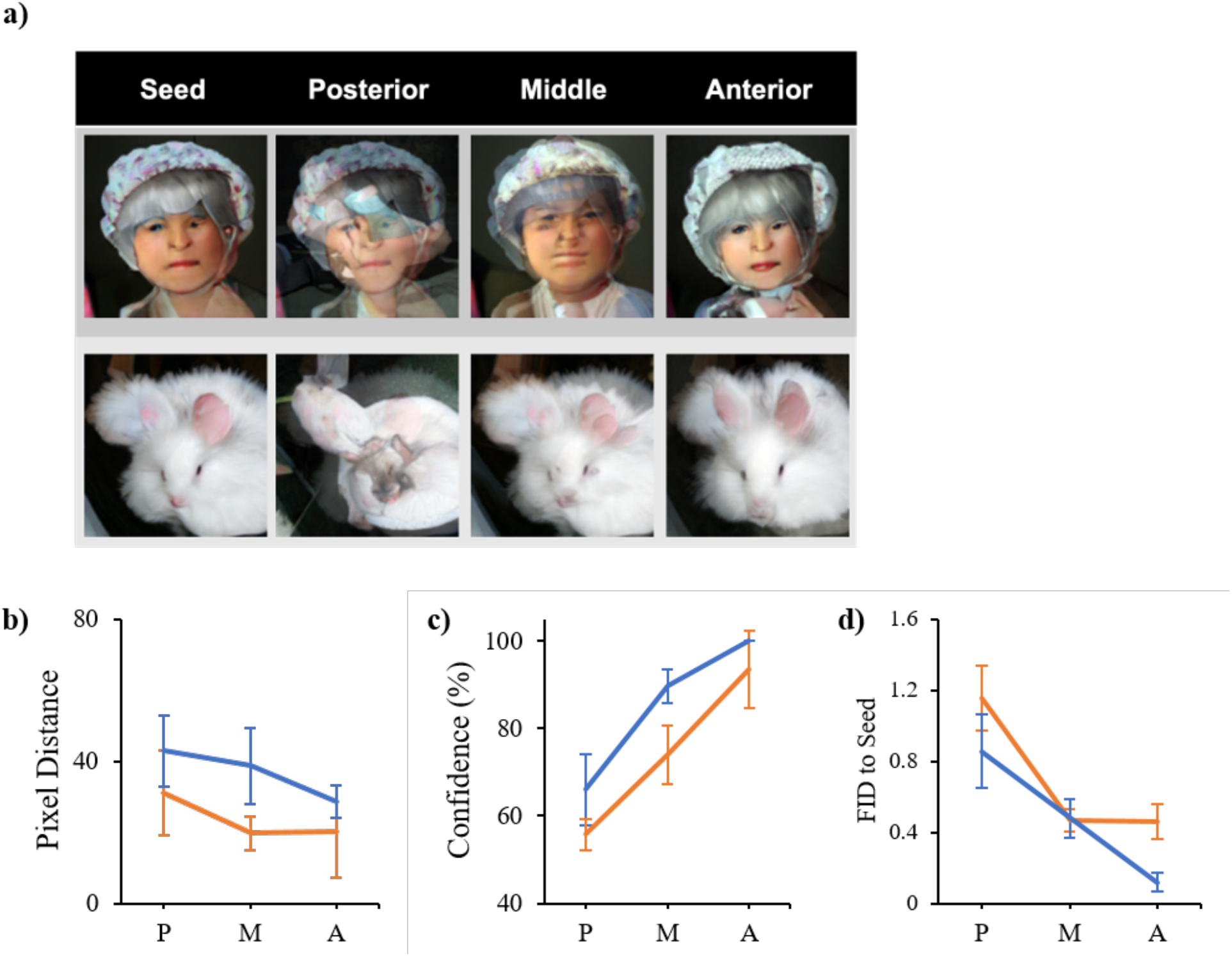
Effects of cortical position on perceptograms. **a)** Examples of perceptograms obtained along the posterior-anterior axis of the central IT cortex. **b)** Pixel distance of seed to the perceptogram. The abscissa represents the position of an LED relative to the posterior-anterior anatomical axis of the central IT cortex. The ordinate shows the pixel distance of the perceptograms resulting from each AP position from one seed image. Blue: Sp, Orange: Ph. While all perceptograms show pixel distance from their seed images, the effect does not change across cortical AP positions on this measure, and the line graph is statistically flat (df = 8 and 11, p = 0.202 and 0.197 for Sp and Ph, respectively, ANOVA). **c)** Classification confidence of a Yolo (real-time object detection system) fed by perceptograms obtained from different cortical positions on the posterior-anterior axis. The abscissa is the same as in b, and the ordinate shows classification confidence. Blue: Sp, Orange: Ph. Classification confidence significantly increases for the perceptograms obtained from anterior cortical positions (df = 8 and 11, p = 0.026 and 0.005 for Sp and Ph, respectively, ANOVA). **d)** FID distance from the seed. The abscissa is the same as in b and c. The ordinate shows the FID of the perceptograms obtained from each LED to its seed image. Blue: Sp, Orange: Ph. FID, normalized to mean, is significantly different across cortical positions (df = 8 and 11, p = 0.009 and 0.011 for Sp and Ph, respectively, ANOVA). Error bars indicate ±1 standard error of mean for all subplots.

### Conclusion

Constructing a mechanistic theory of visual perception requires the establishment of causal homeomorphism between the neural state, a system measured in units of spikes per second, and the perceptual state, a system measured in psychophysical units (*1, 8*). Making the bridge between the two requires parametric characterization of both. Simultaneous measurement of both in large primate brains poses a serious technical challenge. In this study, we decided to focus on characterizing the perceptual events induced by neural stimulation as it has been a historical and methodological bottleneck. The challenge had two faces, one required reliable high-throughput stimulation in a large brain, and the other demanded custom-tailored artificial intelligence in order to develop effective perceptograms. As for the first one, we chose optogenetics over traditional electrical stimulation as it provides more accurate and more interpretable stimulation capacity given that it does not target axons of passage (*28, 29*), and it is less invasive by being a surface implant (*2*). Furthermore, electrical stimulation is not reliable for the high number of stimulation trials required here (*30*). The second face of the challenge demanded not only searching a very large image space but also mimicking the effect of stimulation well enough to deceive the animals against reward. Ahab controlled the search function (see methods), and DaVinci mastered mimicking images by combining two GAN-generated images to achieve an accurate reconstruction of images outside its original training set (see methods).

Facing this two-faced challenge, perceptography provides pictures that are behaviorally exchangeable with the state of being cortically stimulated. Given the parametric nature of these pictures, we can now provide objective and quantitative evidence of the nature and quality of stimulation-driven visual perceptual effects. This allows measurement of what has long been theoretically defined as the “projective field” of neurons (*31*), here operationally defined as “the causal contribution of a given neural group onto the perceptual space.” Characterization of neural projective fields in the visual system, once combined with descriptions of neural sensory response fields (*22*), establishes the missing link between neural activity and perception. This can be done in the context of quantitative modeling that links the two; while existing theoretical models of visual hallucinations yield surprisingly similar results to our observations (*32, 33*), further research is needed in order to complete the picture completion of these steps will provide access to the building blocks of a potential unifying mechanistic theory of perception and consequently provides a deeper understanding of visual hallucinations in mental disorders. It also allows the development of better visual prosthetic devices that, in light of these results, may consider stimulation of the high-level visual cortex in addition to the traditional primary visual cortex.

Altogether, given that the amount of work left to be done in this important area is practically beyond the working bandwidth of a single lab, we find this adventure incomplete yet mature enough to be shared with the scientific community. We hope this work sparks interest in those interested in underlying mechanisms of visual perception and encourages technique developers to invest in platforms that allow easy high-throughput simultaneous recording and stimulation of the cortex in large brains.

## Methods

### The optical array

Using a custom-made injection array (*34*), we performed multiple injections of AAV5-CamKII-(C1V1)-eYFP in a ∼5×5mm region, covering the lateral bank of the central IT cortex. 10uLit of the virus was injected at each track. We later implanted OptoArrays (Blackrock Neurotech) on the virally transduced area as well as the same anatomical region in the opposite hemisphere not injected with the virus. The 3D models of the animals’ brains and skulls were reconstructed with the FLoRIN method to facilitate the surgery and LED placement (*35, 36*). At each “stimulated” trial, one LED on the array was activated. The LED illumination levels used varied depending on the experiment and location on the cortex, but it was kept between 1.5 and 11 mW of total photometric output, adjusted to keep the animal’s performance below the behavioral ceiling. The choice of LED and illumination power was kept constant at each perceptography cycle. More technical details about the array, as well as more behavioral results, can be found in Azadi et al. (*6*)

### Psychophysics

The experiments were performed in a well-lit test chamber in order to avoid retinal dark adaptation that could potentially help the animals detect the cortically delivered light through their skull (see Azadi et al. for more). The animals sat 57cm away from a calibrated screen (32”, 120Hz, 1920×1080 IPS LCD, Cambridge Research System Ltd). The data was collected using a custom MWorks script (*37*) and a Mac Pro 2020. Eye tracking was performed using an Eyelink 1000 Plus (SR Research). All of the behavioral and surgical procedures used in this study were in accordance with the NIH guidelines.

### DaVinci

DaVinci, our illustrator engine, was built based on BigGAN 18(Brock et al. 2019). In order to construct stimuli, DaVinci superimposed a random image over the seed image (both generated in BigGAN), then randomly perturbed the image parameters as well as the transparency of the top layer. The altered image parameters included image class involvement (out of 1000 classes of ImageNet), truncation factor, and the z vector. Given our preliminary results (see fig 1.d), we figured that most of the image search would happen not too far from the seed image; the two-layered image structure was considered to ease this. Nevertheless, we wanted DaVinci to be capable of venturing far and creating virtually any image by varying image parameters as well as layer transparency. To test this, we created seven target images that were not included in DaVinci’s training set (ImageNet) and forced DaVinci to start from a random image seed and recreate the target image in an iterative process using a pixel dissimilarity loss function. The target images ranged from the picture of the dinner plate of one of the authors to modern art pieces warped in photoshop. In all cases, DaVinci recreated the target image with high fidelity (mean pixel similarity= 17.44%, StD=4.28)(See supplementary Fig S2).

### Ahab

Ahab was the optimizer that logged the behavioral responses and navigated DaVinci in order to find the perceptogram. The Ahab algorithm included a VGG-16 (*38*)convolutional neural network (pretrained on ImageNet) as a feature extractor that kept track of the features of the images that satisfy the following criteria: FA rate > 50% and Miss rate < 5%. By extracting and putting together the most common features from the selected images, Ahab created an image prototype called average-feature-prototype (AFP). Then Ahab created a pool of images sprayed around the AFP in the image space to achieve the range of image parameters in the vicinity of the AFP. Based on these parameters, Ahab guided DaVinci to make 2-6 mutants for each image.

## Acknowledgments

We thank Mark Eldridge and Reza Azadi for their crucial contributions to the surgeries. We thank Reza Azadi for the initial training of monkey Ph. This research was supported by the Intramural Research Program of the NIMH ZIAMH002958 (to A.A.)

## Contributions

Concept: A.A

Experiment design: E.S and A.A

Data collection and analysis: E.S and T.M

Designing Ahab and DaVinci: E.S, M.P, and W.S

Writing the manuscript: A.A, E.S, and T.M

## Data and material availability

The data and material that support the findings of this study will be available online following the publication.

---

**Supplementary Fig 1.**
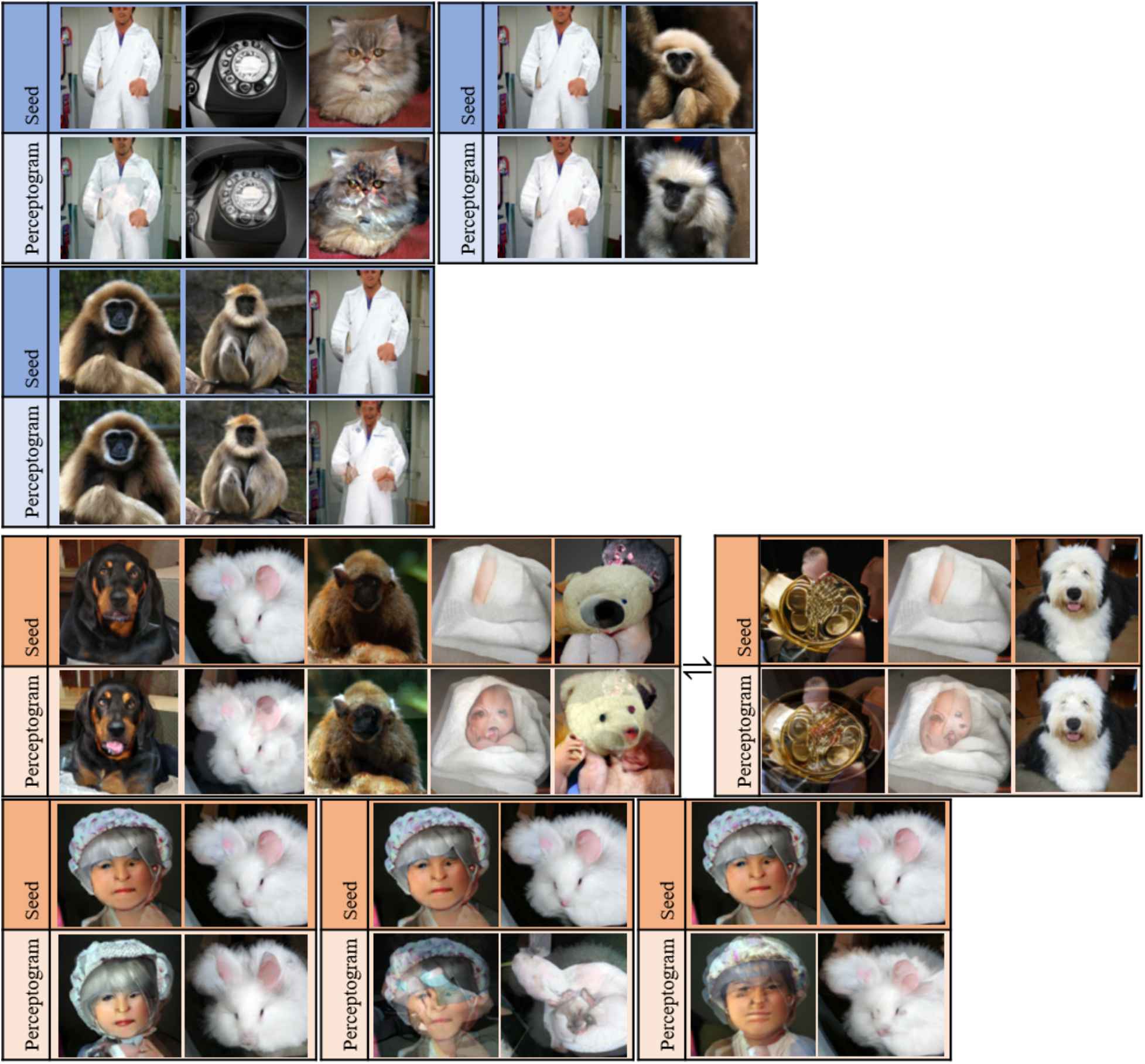
Perceptograms. The rectangular boxes surrounding groups of images indicate the perceptograms obtained from the same cortical position using different seed images for the high intensity stimulation condition. For 3 seed images, in monkey Ph, we tried even higher stimulation intensity. The “⇌” sign shows the blocks from the same cortical position but different stimulation intensities. Please note that subtle changes from the seed image are sometimes hard to see in these small static images, but they are far more visible when the seed image and the perceptogram are viewed in temporal sequences, such as in the experiment. Blue: Sp, Orange: Ph

**Supplementary Fig 2:**
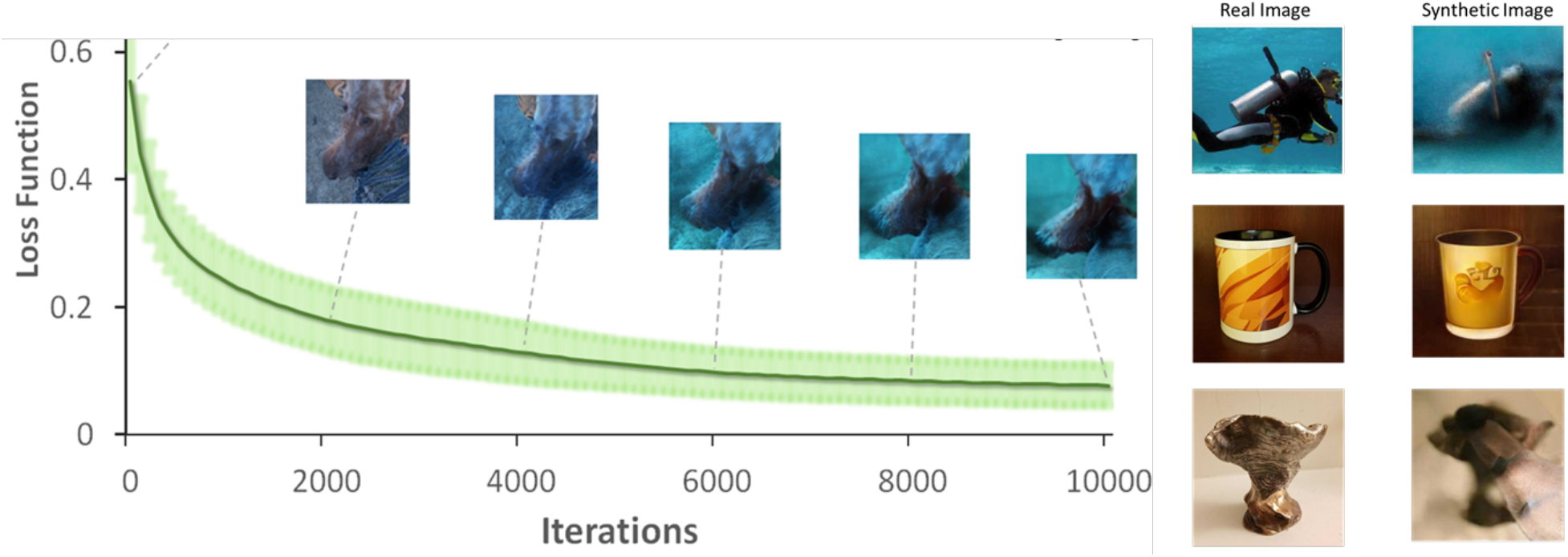
Recreating random images with Ahab and Davinci. To test the quality of Ahab optimization and Davinci image illustration, seven random images (three shown here) \ that were not in the ImageNet (BigGAN and Davinci pretrained sets) dataset were selected. Ahab was then used to optimize a random Davinci image to be as close to the target as possible. The abscissa shows the loss function (pixel distance), and the ordinate represents Ahab iteration cycles. The shaded green shows ±1 standard error of the mean. Examples of images are shown; natural images in the left column and Ahab-optimized images in the right column.

